# The helminth-derived effector protein HpBARI trimerises to selectively target membrane bound ST2 receptor

**DOI:** 10.64898/2026.09.29.755424

**Authors:** Abhishek Jamwal, Nicole W P Ong, Suzanne H Hodge, Henry J McSorley, Matthew K Higgins

## Abstract

The ST2 receptor is activated by the alarmin cytokine IL-33, inducing and controlling immune responses. ST2-mediated signalling contributes to protection from parasitic infection, but is also involved in asthma, inflammatory bowel disease, fibrosis and allergic inflammation, making inhibitors of ST2 desirable. While signalling is mediated by membrane-associated ST2 on immune cells, extracellular fluids also contain a soluble form of ST2 which acts as a decoy, soaking up IL-33. An effective ST2-targeting therapeutic will selectively target membrane-associated ST2. The intestinal helminth *H. polygyrus bakeri* suppresses ST2- mediated signalling to reduce helminth clearance from the gut, with the effector HpBARI2 directly binding to and inhibiting ST2. Here, we present the structure of the ST2:HpBARI2 complex. HpBARI2 forms a stable, planar trimeric structure, with three vertices each binding a single ST2. HpBARI2 occludes the IL-33 binding site of ST2 and constraints the receptor in a conformation incompatible with recruitment of signalling co-receptor IL-1RAcP. The trimeric valency of HpBARI2 results in preferential binding to multiple membrane-associated ST2 molecules rather than the soluble decoy receptor. Therefore, HpBARI2 combines steric blockade, conformational restriction and multivalent binding to potently and selectively inhibit ST2 signalling in the context of soluble ST2. This insight will guide design of improved therapeutics which target ST2.

## Introduction

Activation of immune responses is important for protection from infection but can also cause severe inflammatory conditions when dysregulated (*1, 2*). The interleukin-1 family receptor ST2 (IL-1RL1) is expressed on various immune cells involved in type 2 immune responses, including ILC2 cells (*3, 4*). The alarmin cytokine IL-33 binds the ST2 ectodomain and stabilises it in a conformation that binds the co-receptor IL-1RAcP to form a signalling competent heterodimer (*5, 6*). This induces expression and secretion of cytokines, including IL-5 and IL-13, which amplify or recruit effector cells and mediate a type 2 response (*1*). ST2 signalling can be protective, especially against intestinal helminth infections. However, when it is inappropriately triggered it can cause allergic and fibrotic diseases, including asthma, making ST2 a potential therapeutic target (*1*). Indeed, ST2 inhibitors are currently being investigated for treatment of inflammatory disorders including asthma, chronic obstructive pulmonary disease and intestinal fibrosis (*7–10*). However, serum and other extracellular fluids also contain a soluble form of ST2 (sST2), which acts as decoy receptor by binding to IL-33 and blocking its activity (*11, 12*). Inhibitors of ST2 must function in the context of this soluble ST2 to access signalling competent membrane-associated ST2.

Resistance to intestinal helminths is mediated by type 2 immunity and the IL-33 pathway plays a role, with significantly increased susceptibility in mice deficient in IL-33 or ST2 (*13, 14*). Helminths such as the murine intestinal parasite *Heligmosomoides polygyrus bakeri* have evolved a cocktail of secreted immunomodulatory products, including regulators of type 2 immunity and of the IL-33 pathway (*15, 16*). Amongst these excretory-secretory products are two distinct proteins families, each consisting of complement control protein (CCP) domains. The HpARI family bind IL-33 while the HpBARI family bind ST2, thereby targeting both sides of the IL-33:ST2 signalling complex (*17, 18*). Members of the HpARI family show differential activity against IL-33 from different host species, depending on whether or not they block IL-33 from binding to ST2 (*17, 19–22*). The two HpBARI proteins, HpBARI and HpBARI_Hom2 (hereafter HpBARI2), both suppress type 2 immunity by interacting with the ST2 ectodomain, although their mechanism of action is not yet understood (*18*).

HpBARI2 is of particular interest as it can bind both human as well as murine ST2, preventing immune cells from responding to IL-33 mediated activation (*18*). In addition, in mouse models, injection with HpBARI2 reduced allergic airway inflammation, promoted bone and joint repair in haemophilic arthropathy, and improved survival during SARS-CoV-2 infection, indicating multiple potentially valuable therapeutic outcomes of HpBARI2-mediated inhibition of ST2 signalling (*18, 23, 24*). These observations identify HpBARI2 as a potential therapeutic inhibitor of ST2 signalling and raise the question of how this soluble effector achieves effective inhibition of ST2 signalling in the context of sST2. Here, we combine structural and biophysics studies with cellular signalling assays to reveal how HpBARI2 selectively and effectively blocks membrane-bound ST2, guiding future development of ST2 inhibitors.

## Results

### HpBARI2 forms a trimeric assembly that engages three ST2 receptors

We first aimed to determine the structure of HpBARI2 bound to the *Mus musculus* ST2 receptor ectodomain (ST2^ecto^). We expressed both components in HEK293F cells, purified them and mixed to form a complex. Size exclusion chromatography (SEC) analysis, with and without multiangle laser light scattering (SEC-MALLS), showed that ST2^ecto^ and HpBARI2 form a homogenous ∼173 kDa complex, which is around three-fold larger than expected for a complex of a single HpBARI2 bound to a single ST2 ectodomain (Fig. S1A). The size of the complex made it suitable for structure determination using cryogenic electron microscopy and single particle analysis (Fig. S1B). The structure revealed a planar HpBARI2 trimer which engages three ST2^ecto^ molecules, one at each vertex of the triangle (Fig. 1A).

**Fig. 1.**
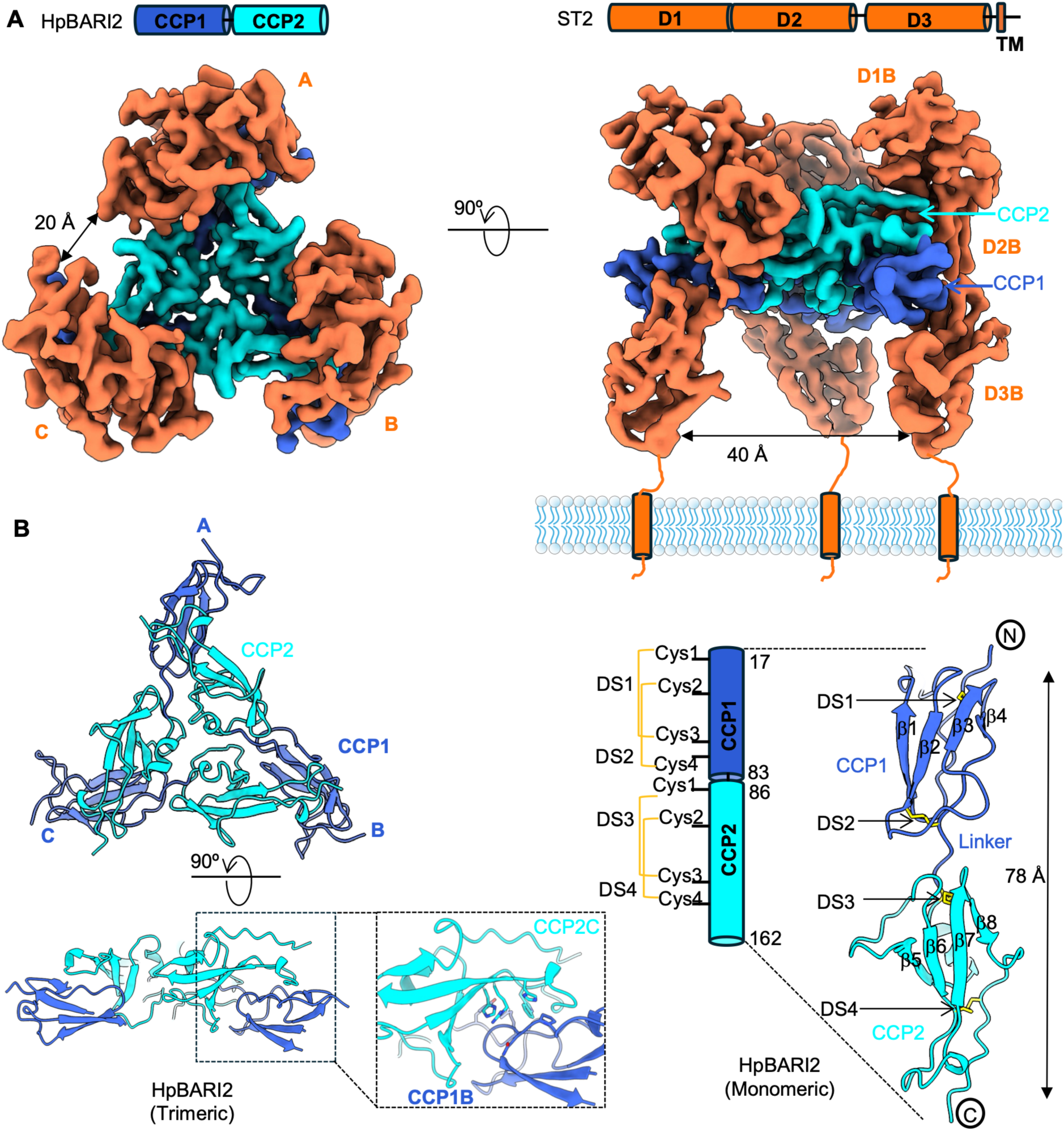
Structure of the ST2^ecto^:HpBARI2 complex. **(A)** Two views of the sharpened cryogenic electron microscopy-derived map of the ST2^ecto^:HpBARI2 complex with three ST2 molecules (coral) binding to HpBARI2 trimer. The CCP1 and CCP2 domains of HpBARI2 are shown in blue and cyan respectively. Above the structure are shown domain diagrams of the primary structures of HpBARI2 and ST2. The three ecto-domains (D1–D3) of ST2 are shown together with the transmembrane region (TM). **(B)** Cartoon representations of trimeric and monomeric HpBARI2. The left panel shows three extended HpBARI2 monomers forming a trimeric structure in which CCP2 domains from each subunit form a trimeric core. At each vertex the CCP1 and CCP2 domains from neighbouring subunits pack together. The enlarged view in the dotted box shows the CCP1:CCP2 interface with selected interacting side-chain residues shown as sticks. The right- hand panel shows a single HpBARI2 subunit, with secondary structure elements and disulphide bonds (DS1-4) indicated.

Each HpBARI2 subunit comprises two CCP domains (CCP1 and CCP2) connected by a two-residue linker (Fig. 1B). Extensive interactions involving both domains mediate formation of a stable trimer. The three CCP2 domains are located at the core of the trimer and collectively bury ∼ 645 Å^2^ through inter-subunit contacts. At each vertex, the CCP1 domain of one subunit packs against loops emerging from the C-terminal end of CCP2 of the neighbouring subunit to bury ∼ 800 Å^2^ per interface (Fig. 1B). The HpBARI2 trimer forms three binding sites for ST2^ecto^, with one binding each vertex (Fig. 1A). These sites are independent, with the three receptors separated by ∼20 Å at the membrane distal side and ∼40 Å at the membrane proximal side (Fig. 1A). The concave face of each ST2^ecto^ wraps around its binding site on HpBARI2, with all three domains of ST2 contacting HpBARI2.

Each binding site contains residues contributed from CCP1 of one HpBARI2 subunit and from CCP2 of another subunit, indicating that trimerization is required for ST2 engagement. To test this, we introduced a G26R mutation in CCP1, designed to disrupt inter-subunit packing of CCP1 and CCP2 by steric hinderance (Fig. S2A). SEC-MALLS showed that HpBARI2^G26R^ has a molecular weight of ∼18 kDa, compared with 59 kDa for HpBARI2, consistent with disruption of the trimer into monomeric species (Fig. S2B and C). We next tested binding of HpBARI2 and HpBARI2^G26R^ to immobilised ST2^ecto^ by surface plasmon resonance (SPR). HpBARI2 produced a clear and robust binding response at 5nM concentration, while 500nM HpBARI2^G26R^ showed no response (Fig. S2D). Therefore, disrupting the HpBARI2 trimer abolished ST2 binding, confirming that assembly of trimeric HpBARI2 is required for ST2 binding.

### HpBAR2 blocks ST2 through steric competition and conformational constraint

We next aimed to characterise how ST2 is inhibited by HpBARI2. We first designed a mutant HpBARI2 which lacks ST2 binding. Analysis of the structure reveals that three distinct interfaces stabilize the ST2^ecto^:HpBARI2 complex (Fig. S3A). Interface 1 consists primarily of electrostatic interactions between the CCP2 domain of HpBARI2 and the D1-D2 domains of ST2^ecto^. Addition of a C-terminal His_6_ tag to HpBARI2 was predicted to disrupt this interface and reduced binding by 20% and increasing the dissociation rate (Fig. S3B). Interface 2 involves hydrophobic and polar interactions between the β1-β3 surface of CCP1 and the D3 domain of the receptor (Fig. S3A) and the V66R mutant sterically occluded this interface and reduced ST2^ecto^ binding by ∼70% (Fig. S3C). Interface 3 is formed by the opposite face of CCP2 from interface 1. This engages with the convex side of domain D2 of ST2^ecto^ (Fig. S3A). Y141R and L146R mutations were predicted to disrupt interface 3 and together reduced binding by ∼50% (Fig. 2A and Fig. S3D). While disruption of any individual interface did not abolish binding, perhaps because of the flexibility of ST2 and the complexity of its binding to HpBARI2, combining three sets of interface disrupting mutations into one HpBARI2 completely abolished ST2^ecto^ binding (Fig. 2A and Fig. S3E and F).

**Fig. 2.**
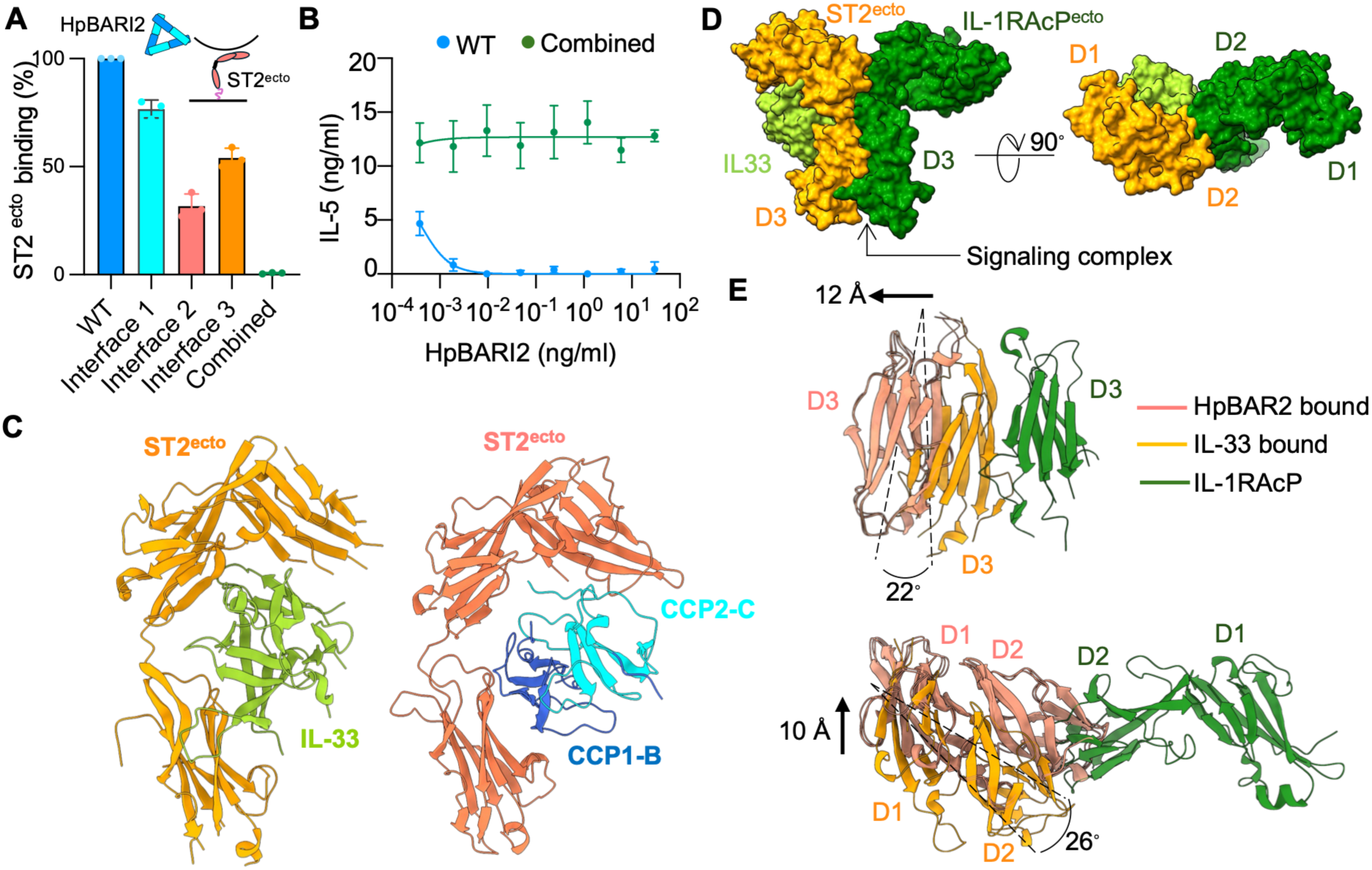
Binding and inhibition of ST2 signalling by HpBARI2. **(A)** Binding of wild-type HpBARI2 and mutants with disrupted ST2 interfaces to immobilized ST2^ecto^, as determined by SPR at a HpBARI2 concentration of 5 nM. Combined as mutations in all three interfaces. Bars represent the mean ± standard error of mean (SEM) of three independent measurements (n = 3). Schematic shows the experimental setup. **(B)** Concentration-response curves showing inhibition of ST2 signalling by HpBARI2 and the mutant of HpBARI2 with all three interfaces disrupted (combined). Error bars represent the mean ± standard error of the mean (SEM) of three biological replicates. Data are representative of two independent experiments (n = 2). **(C)** Cartoon representation showing a side-by-side comparison of ST2^ecto^ (light orange) bound to IL-33 (green) (left-hand panel, PDB ID: 5VI4) with ST2^ecto^ (coral) bound to HpBARI2 (blue and cyan) (right-hand panel). **(D)** Surface representation of the ternary signalling complex between ST2 (orange), IL-33 (green) and IL-1RAcP (green). **(E)** Cartoon overlays showing the domain rearrangements in HpBAR2- bound ST2^ecto^ (coral) relative to IL-33-bound ST2^ecto^ (orange) from the ternary signalling complex. Upper panel: The D1-D2 domains of ST2^ecto^ from the HpBARI2 bound complex were superimposed onto the corresponding D1-D2 domains of ST2^ecto^ in the ternary complex, revealing the position of D3 domains in ST2^ecto^:HpBARI2 complex. Lower panel: The D3 domain of ST2^ecto^ from HpBARI2 bound complex was superimposed onto the corresponding D3 domains of ST2^ecto^ in the ternary signaling complex, revealing the resulting position of D1-D2 domains in the ST2^ecto^:HpBARI2 complex.

We next tested whether disruption of these three interfaces affects the ability of HpBARI2 to inhibit ST2 signalling. We used a murine bone marrow-derived ILC2 assay in which IL-5 secretion is measured as an output of IL-33-mediated ST2 signalling (*19*). In this assay, wild-type HpBARI2 suppressed IL-5 secretion even at pg/ml concentrations, whereas the combined interface mutant which failed to bind ST2^ecto^ had no detectable suppressive activity at 10,000-fold higher concentrations (Fig. 2B). These results confirm that the structural interfaces identified in the cryo-EM complex are necessary for HpBARI2-mediated inhibition of ST2 signalling.

We next investigated how HpBARI2 blocks ST2 signalling by comparing our structure of HpBARI2-bound ST2^ecto^ with a previous structure of ST2 bound to both IL-33 and the IL- 1RAcP co-receptor (*5*). ST2^ecto^ adopts a similar structure when bound to both IL-33 and HpBARI2, with the two ligands sharing overlapping interfaces on ST2^ecto^ (Fig. 2C). Therefore, HpBARI2 directly and sterically occludes the binding site for IL-33, consistent with our previous observation that ST2^ecto^ pre-bound to HpBARI2 cannot bind IL-33 (*18*).

ST2 signalling also requires recruitment of the co-receptor IL-1RAcP, raising the possibility that HpBARI2 binding may also interfere with formation of the IL-33:ST2:IL-1RAcP signalling complex (Fig. 2D). We tested this by SPR using immobilized IL-1RAcP. While the ST2^ecto^:IL-33 complex bound IL-1RAcP, the ST2^ecto^:HpBARI2 complex showed no detectable binding (Fig. S4A). In addition, SEC analysis showed that IL-1RAcP formed a ternary complex with ST2^ecto^ and IL-33 but did not co-elute with ST2^ecto^:HpBARI2 (Fig. S4B and C). Thus, HpBARI2 binding not only competes with IL-33 but also prevents co-receptor engagement.

To reveal how HpBARI2 stops ST2 from binding IL-1RAcP, we overlaid the structure of ST2^ecto^ bound to HpBARI2 onto the crystal structure of the ST2^ecto^:IL-33:IL-1RAcP^ecto^ ternary complex (*5*). This revealed that HpBARI2 and IL-33 stabilise different conformations of ST2^ecto^. In a particular, there are differences in the relative positions of D1 and D2 due to a ∼26° rotation, creating a ∼10 Å displacement of D2 relative to D1. There is also a difference in the rotational position of the D3 domain by ∼22° together with a ∼12 Å displacement. This stabilises ST2 in a conformation incompatible with binding IL-1RAcP (Fig. 2E). Together, our findings support a mechanism which combines steric occlusion and conformational constraint as mechanistic basis of ST2 antagonism by HpBARI2.

### Trimerisation of HpBARI2 allows improved binding to membrane-associated ST2

Having established that trimeric HpBARI2 binds three ST2^ecto^ molecules, we next asked whether multivalent binding provides avidity to allow selective binding to membrane-associated ST2. We reasoned that trimeric HpBARI2 will simultaneously bind to three membrane-associated ST2 molecules with the resultant avidity stabilising binding, without changing the affinity for monomeric soluble ST2. To test this, we designed a monovalent HpBARI2 heterodimer (HpBARI2^mono^) that contains a single functional ST2 binding site (Fig. 3A). This was produced by co-expressing two HpBARI2 variants carrying three distinct but harmonising sets of mutations: i) mutations that disrupt CCP1:CCP2 inter-subunit packing; ii) mutations to create a stabilising disulphide bond and iii) an interface mutation and a modification that eliminates residual ST2 binding to disrupted CCP domains (Fig. S5A).

**Fig. 3.**
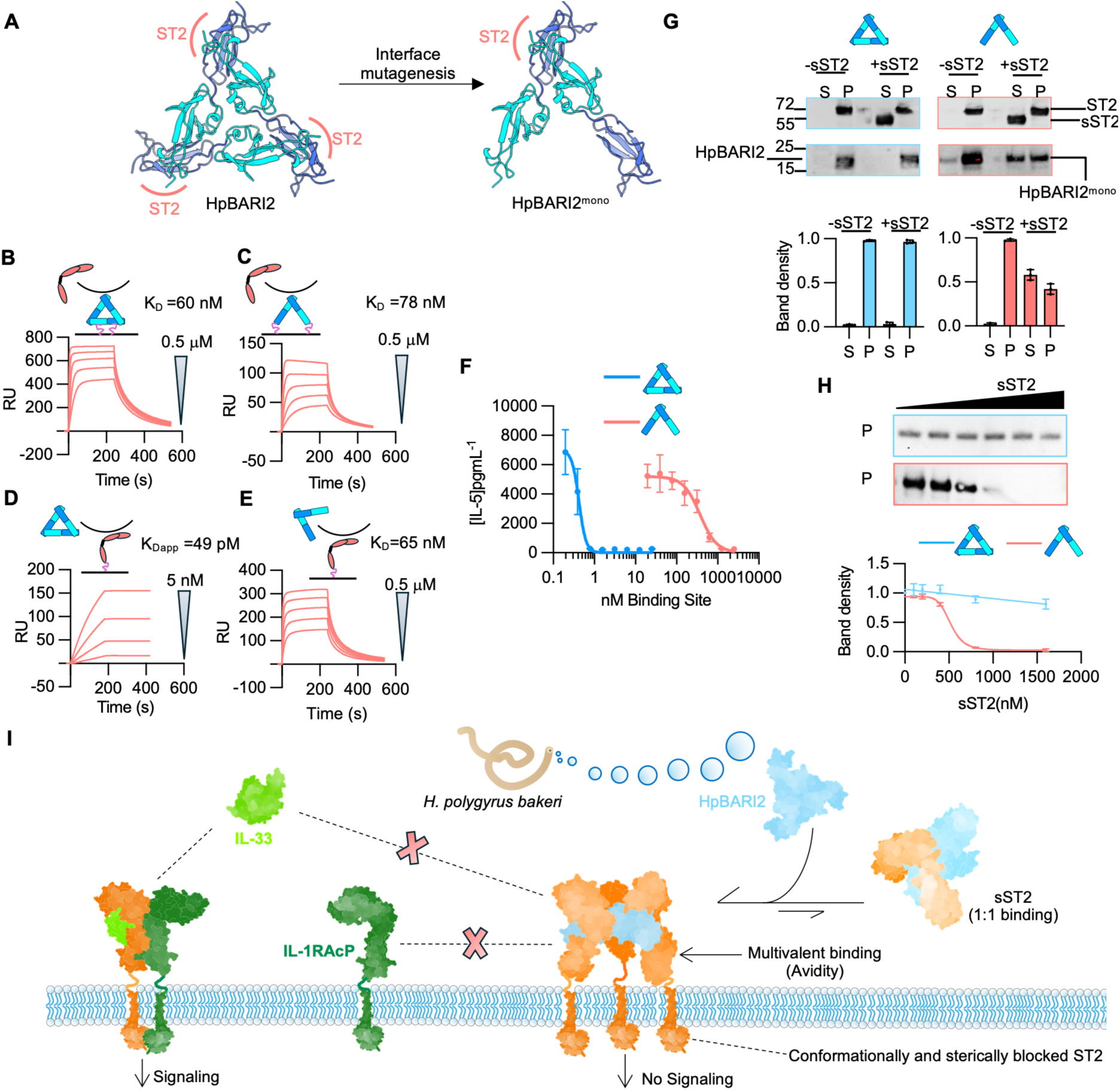
HpBARI2 preferentially binds membrane-bound ST2 through avidity. **(A)** Cartoon representations illustrating trimeric HpBARI2 (left) and a monovalent heterodimeric form, HpBARI2^mono^) (right). **(B-E)** SPR analysis of HpBARI2 and HpBARI2^mono^ binding to ST2^ecto^ using two experimental formats. Schematics show the experimental formats. In the first format, biotinylated HpBARI2 (**B**) or HpBARI2^mono^ (**C**) were immobilized on the chip surface, and binding responses were measured by flowing two-fold serial dilutions of ST2^ecto^, starting at 500 nM. In the second format, ST2^ecto^ was immobilized, and binding responses were measured by flowing two-fold serial dilutions of HpBARI2 (**D**) or HpBARI2^mono^ (**E**), starting at 500 nM and 5 nM, respectively. Apparent K_D_ values were determined by Langmuir fitting and represent two independent experiments (n = 2). **(F)** Concentration response curves showing inhibition of signalling by HpBARI2 and HpBARI2^mono^. Error bars represent the mean ± standard error of the mean (SEM) of three technical replicates. **(G)** Western blot analysis (top panel) showing how soluble ST2 (sST2) affects binding of HpBARI2 or HpBARI2^mono^ to liposome-embedded ST2. Molecular masses are indicated in kDa. The accompanying plots (bottom panels) show band densities of HpBARI2 or HpBARI2^mono^ in the supernatant and pellet fractions. Error bars represent the mean ± SEM of three technical replicates. **(H)** Western blot analysis (top) and band densitometry (bottom) showing the effect of a two-fold serial dilution of sST2 (1000–61.25 nM) on HpBARI2 or HpBARI2^mono^ binding to membrane-embedded ST2. Error bars represent the mean ± SEM of three technical replicates. **(I)** Proposed model for HpBARI2 function. Multivalent receptor engagement allows avidity and selective antagonism of membrane- bound ST2, even in the presence of soluble ST2. This directly blocks access of IL-33 and IL- 1RAcP to the signalling receptor.

Coomassie stained SDS-PAGE gels showed that, under nonreducing conditions, the purified protein migrated at nearly twice its molecular weight, suggesting formation of a disulphide-linked heterodimer (Fig. S5B) and SEC-MALLS analysis revealed a complex of ∼78 kDa, consistent with one HpBARI2^mono^ (42 kDa) bound to one ST2^ecto^ (38 kDa) (Fig. S5D).

We next compared the kinetics of ST2 binding by HpBARI2 and HpBARI2^mono^. We first immobilised HpBARI2 and HpBARI2^mono^ on the sensor chip and flowed over ST2^ecto^, selecting this configuration to ensure that we measure monovalent affinities. Both HpBARI2 and HpBARI2^mono^ exhibited rapid association and dissociation kinetics with similar apparent affinities of ∼60 nM for HpBARI2 and ∼78 nM for HpBARI2^mono^, showing that the single ST2- binding site on HpBARI2^mono^ is correctly structured (Fig. 3B and C). We next assessed binding of HpBARI2 and HpBARI2^mono^ to immobilized ST2^ecto^, allowing avidity through binding of one HpBARI2 trimer to multiple ST2. In this case, the binding profiles for HpBARI2 and HpBARI2^mono^ differed substantially (Fig. 3D and E). HpBARI2^mono^ displayed rapid dissociation with apparent affinity of 65 nM, matching the previous monovalent affinity. In contrast, HpBARI2 bound more slowly with a slow off-rate and an apparent affinity of ∼ 49 pM. When soluble ST2^ecto^ was flowed over the HpBARI2 coated ST2^ecto^ surface, no further binding was observed, suggesting saturation of the ST2-coated chip with HpBARI2 (Fig. S6A). This confirms that trimerization of HpBARI2 allows it to simultaneously engage multiple ST2 molecules, with this avidity leading to very strong binding to immobilised ST2.

We next determined whether avidity increases the ability of HpBARI2 to block ST2 signalling. Murine bone marrow-derived ILC2 cells were incubated with the same molar concentration of HpBARI2 and HpBARI2^mono^. While HpBARI2 inhibited IL-5 secretion with IC_50_ ∼0.3 nM, ∼1,000-fold more HpBARI2^mono^ was required to generate the same effect, with IC_50_ ∼300 nM (Fig. 3F). Therefore, trimerization of HpBARI2 increases its efficacy in blocking ST2-mediated signalling through avidity.

### Trimerisation of HpBARI2 allows selective binding to membrane-associated ST2 in the presence of soluble ST2

Finally, we hypothesized that avidity could also enable HpBARI2 to preferentially bind membrane-embedded ST2 in the presence of sST2. This would be beneficial to the helminth as it would allow HpBARI2 to selectively bind and inhibit membrane-bound ST2 even in the presence of high concentrations of the non-signalling sST2 decoy receptor. To test this, we expressed and purified full length ST2 and reconstituted it into liposomes (ST2somes) (Fig. S6B). We then assessed the binding of HpBARI2 and HpBARI2^mono^ to ST2somes in the presence and absence of sST2 (Fig. S6B). Western blot analysis showed that HpBARI2 and HpBARI2^mono^ co-pellet with ST2some in the absence of sST2 (Fig. 3G and Fig. S6C). Addition of sST2 had little effect on HpBARI2, with most of the protein still found in the ST2somes pellet (Fig. 3G). However, sST2 substantially inhibited association of HpBARI2^mono^ with ST2somes, leaving only ∼40% of HpBARI2^mono^ in the pellet fraction (Fig. 3G). Increasing sST2 concentrations increasingly inhibited HpBARI2^mono^ binding to ST2somes, whereas HpBARI2 remained relatively resistant to competition (Fig. 3H and Fig. S6D).

Together, these findings show that HpBARI2 trimerization allows multivalent binding to membrane-associated ST2. This decreases the amount of HpBARI2 required to inhibit ST2 signalling and drives preferential targeting of membrane bound ST2 over sST2. This avidity driven mechanism is likely to allow HpBARI2 to selectively suppress ST2-mediated signalling, even in the presence of high concentrations of the sST2 decoy during *H. polygrus bakeri* infection (Fig. 3I).

## Discussion

Inhibitors of IL-33-mediated signalling are in active development, with the aim of disrupting signalling processes which are triggered in diseases including asthma and allergy (*1, 25*).

Indeed, injection of Astegolimab, an anti-ST2 antibody, resulted in a modest reduction in asthma exacerbations in a phase IIb study (*7*). However, the anti-ST2 antibody GSK3772847 (melrilimab) lowered soluble ST2/IL-33 concentrations but did not significantly improve clinical symptoms in asthma or eczema, leading to a pause in development (*26, 27*). These findings suggest that, while inhibiting ST2 does have merit, more insight is required to design highly effective biologics which block ST2 function in a physiological context. This raises the question of what we can learn from HpBARI2 about the mechanism of action of highly effective helminth-derived modulators of ST2 signaling which we can apply to design of novel biologics.

The mechanism by which IL-33 mediates signalling has been characterised through structural and mechanistic studies of the IL-33:ST2:IL-1RAcP complex (*5*). However, little was known about how ST2-mediated signalling can be effectively inhibited. Here, we show how the helminth effector HpBARI2 inhibits ST2. We show that HpBARI2 sterically blocks the IL- 33 binding site on ST2 and stabilises a conformation of ST2 incompatible with forming the IL-33:ST2:IL-1RAcP complex. This is conceptually similar to the function of canonical IL-1 family receptor antagonists IL-1Ra and IL-36Ra. These cytokine analogs occupy their cognate receptors while preventing conformational changes needed for signalling (*28–30*). However, a major difference is that HpBARI2 adopts a trimeric architecture, allowing it to simultaneously bind to three ST2 molecules. This increases the stability of binding of HpBARI2 to membrane-associated ST2 and reduces the concentration required to inhibit signalling by ∼1000-fold, with avidity substantially increasing the potency of HpBARI2.

The discovery that HpBARI2 uses avidity to selectively bind a membrane-associated receptor may have broader implications for inhibitor design. IL-33-mediated signalling occurs in the presence of soluble ST2. This acts as a decoy receptor which can sequester IL-33 before it reaches ST2 receptors at immune cell surfaces. Soluble ST2 is also likely to provide a challenge for therapeutic agents which target ST2-mediated signalling, increasing the active concentration required for efficacy by sequestering the agent before it reaches its site of action. This is not unique to ST2, as soluble forms of other immune receptors, including those for IL-1 (34) and TNF (35), have also been detected. HpBARI2 has solved this challenge by evolving only moderate monovalent affinity while achieving high potency instead through multivalent engagement with membrane associated receptors. Indeed, an increased monovalent affinity would equally increase the binding to decoy receptors. Our findings therefore raise the possibility that maximizing monovalent affinity may not always be optimal for receptor antagonists which function in the context of soluble decoy receptors. Instead, biologics with moderate or weaker affinity featuring rapid dissociation, which generate potency through multimerisation, can preferentially target membrane receptors while limiting neutralization by soluble receptors. Such an approach may be relevant in inflammatory settings where soluble receptors are present (*31–33*).

Our study therefore shows that HpBARI2 blocks ST2-mediated signalling by occluding IL-33 binding and preventing ST2 from adopting a conformation compatible with formation of the signalling complex. It also uses multivalent binding to selectively inhibit ST2 in the context of soluble ST2 decoy receptor. This study shows how a naturally evolved parasite effector has evolved to target ST2-mediated signalling and can inform the design of next-generation ST2- targeting biologics.

## Methods

### Protein expression and purification

#### HpBARI, ST2^ecto,^ IL-1RAcP^ecto^

Coding DNA sequences corresponding to HpBARI2 and *Mus musculus* ST2 ectodomains (ST2^ecto^) were cloned in the pSECTAG-2A vector (Thermofisher) with N-terminal (HpBARI2) and C-terminal (ST2^ecto^) His6 tags for protein production in mammalian expression systems. ExpiHEK293F cells (Thermo Fisher) were transfected with expression constructs using the Expifectamine™ 293 Transfection Kit (Thermo Fisher). Enhancers 1 and 2 were mixed and added to the transfected cell mix 18-24 hours post-transfection. Cell supernatants were harvested on the fourth day post-transfection and filtered using 0.4-micron assembly filters (Nalgene). Ni-NTA Sepharose excel resin (Thermo Fisher) was washed and equilibrated with a wash buffer containing 20 mM HEPES-Na, pH 7.2, 200 mM NaCl and 20 mM imidazole before incubation with filtered supernatants. Following binding, the resin was washed with 10-column volumes of wash buffer. Proteins were eluted using wash buffer containing 500 mM imidazole. To remove misfolded protein aggregates, affinity-purified proteins were concentrated to 3 mg/ml and subjected to size exclusion chromatography (SEC) with HpBARI2 purified using a S75 10/300 increase (cytiva) and ST2^ecto^ using a S200 10/300 increase (cytiva) columns. In both cases, PBS was used. All mutants of HpBARI2 and C-terminal Avi-tagged ST2^ecto^ and *Mus musculus* IL-1RAcP^ecto^ were expressed and purified using the same procedure.

#### Full-length ST2 and sST2

Full-length ST2 was cloned into a pFastBac1 (Thermo Fisher) vector carrying a C-terminal His_6_ tag for expression in Sf9 cells. Bacmids were generated in DH10Bac cells (Life Technologies) and purified by isopropanol precipitation using a miniprep kit (QIAGEN). First- generation baculoviruses (P1) were produced by transfecting Sf9 cells with individual bacmids at a cell density of 1×10^6^ cells/ml and subsequently amplified through two steps to produce third-generation viruses (P3). ST2 expression was induced by adding viral particles from P3 to Sf9 cells (∼3.0-4.0 × 10^6^ cells/ml) at a final concentration of 1.5%. After 48 hours cells were harvested and resuspended in 5mM Tris-Cl, pH 8.0 and lysed using continuous flow cell disruptor (Constant Systems). Lysed homogenate was centrifuged at 3000 × *g* for 20 min to remove cell debris, and the supernatant was spun at 100,000 × *g* for 45 min at 4°C to isolate Sf9 plasma membranes. Isolated membranes were homogenised in lysis buffer (20 mM Tris-Cl pH 8.0, 300 mM NaCl, 10% glycerol, 10 mM imidazole) and solubilised with 1.2% DDM:CHS (10:1) at 4°C overnight. After centrifugation at 100,000 × *g* for 45 min, the supernatant was incubated with Ni-NTA resin (QIAGEN) at 4°C for 2 hr with gentle stirring.

The resin was then packed in a gravity flow column and washed with a buffer containing 25 mM Tris, pH 8.0, 300 mM NaCl, 10% glycerol, 15 mM imidazole, and 0.02% DDM:CHS. The protein was eluted in lysis buffer supplemented with 500 mM imidazole. Eluted protein was concentrated and injected onto a Superdex 200 increase 10/300 GL column (cytiva).

Fractions containing ST2 were pooled and re-purified via Ni-NTA chromatography. Eluted protein was concentrated and further purified via SEC to obtain pure protein with 0.15-0.2 mg full length ST2 obtained from 2L of Sf9 cell culture.

For soluble ST2 or sST2 production, the sST2 coding DNA sequence was cloned into the pFastBac1 vector (Thermo Fisher) containing honeybee melittin signal peptide sequence for production of protein in a soluble secreted form. The virus was produced and the protein expressed using the same strategy as for ST2 expression. Sixty hours post-infection with the P3 virus, the Sf9 cell culture supernatant was harvested, and protein was purified via Ni-NTA chromatography. Cell culture supernatant was incubated with Ni-NTA resin pre-equilibrated with a wash buffer containing 20 mM Tris-Cl pH 8.0, 300 mM NaCl, 10% glycerol and 20 mM imidazole for 30 mins. Ni-NTA resin containing bound protein was packed on a gravity flow column and washed with 10 column volumes of wash buffer. Bound sST2 was then eluted with buffer containing 20 mM Tris-Cl, 150 mM NaCl and 500 mM imidazole. The eluted protein was concentrated and injected into S200 10/300 increase column (cytiva) equilibrated with 1X PBS to obtain aggregate free sST2.

#### Production of a monovalent form of HpBARI2

Monovalent HpBARI2 (HpBARI2^mono^) heterodimer was produced by mixing HpBARI2^G26R/E108C/V66R^ and HpBARI2^W122R/E108C/C6X-His^ constructs in 1:1 ratio and transfecting the mix into ExpiHEK293F cells using expifectamine transfection kit (Thermo Fisher Scientific). Cells were harvested 96 hours post transfection and expressed proteins were purified via Ni-NTA chromatography. Cell culture supernatant was incubated with Ni-NTA resin pre-equilibrated with wash buffer containing 20 mM Tris-Cl pH 8.0, 300 mM NaCl, 10% glycerol and 20 mM imidazole for 30 mins. Ni-NTA resin containing bound protein was packed in a gravity flow column and was washed with wash buffer. Bound HpBARI2^mono^ was then eluted with buffer containing 20 mM Tris-Cl, 150 mM NaCl and 500 mM imidazole. The eluted protein was concentrated and injected into a S75 10/300 increase column (cytiva) equilibrated with 1x PBS to obtain aggregate free HpBARI2^mono^. 5 μl of each SEC fraction was mixed with 1.5 μl of 4x loading dye containing 10 mM or no dithiothreitol (DTT) and separated on SDS-PAGE. Coomassie stained fractions in non-reducing gel lane fractions corresponding to disulphide cross-linked protein were pooled together to obtain a fraction enriched in disulphide cross-linked HpBARI2^mono^. This HpBARI2^mono^ enriched fraction was concentrated and injected into S75 10/300 increase column (cytiva) pre-equilibrated with 1x PBS. SEC fractions were again assessed on Coomassie stained gels under reducing and non- reducing conditions and the cross-linked fractions were pooled and used for downstream experiments. Following these steps, 0.15-0.2 mg of HpBARI2^mono^ was obtained from 100ml of Expi293F cell culture supernatant.

### Structure determination using cryogenic electron microscopy

#### Grid preparation and data collection

To generate protein complex, 100 µg of HpBARI2 was mixed with 500 µg of ST2 and incubated on ice for 30 min. Protein was injected into S200 10/300 increase column (cytiva pre-equilibrated with 1x PBS, to isolate ST2^ecto^:HpBARI2 complex. The obtained complex was concentrated to 4 mg/ml for cryo-EM sample preparation. For cryo-EM, ultrafoil R1.2/1.3 300 Au grids were first glow discharged for 30 secs at 10 mA. Grids were then loaded on a Vitrobot Mark IV (Thermofisher) configured to exert a blot force 0 for 3 seconds. 3.5 µl of protein was applied on the grid and immediately plunge frozen in liquid ethane at 100% humidity. Grids were imaged using an FEI Titan Krios operating at 300 kV and equipped with a Gatan BioQuantum energy filter (20 eV) and K3 direct electron detector. Data collection was automated using fast acquisition mode in EPU (Thermo Fisher). Images were acquired at a nominal 58,149-fold magnification corresponding to a calibrated pixel size of 0.832 Å, at a dose rate of 16.32 electrons Å^−2^ s^−1^ and total exposure time of 3 s with 40 frames. This resulted in a total dose of 48.97 electrons Å^−2^. Images were acquired using a 100 µm objective aperture with defocus range of -3.0 to -1.0 µm in 0.25 µm increments. A total of 4893 movies were acquired.

#### Image processing

Image processing was performed using cryoSPARC (*34*). Movies were motion corrected using patch motion correction, followed by patch contrast transfer function estimation on patch motion corrected micrographs. Initial particle picking was performed using blob picker on 500 micrographs. Picked particles were extracted with a box size of 240 pixels. After two rounds of 2D classification, particles corresponding to eleven representative 2D classes were used as templates for particle picking on all micrographs. Following this procedure, 2,037,659 particles were extracted and subjected to four rounds of 2D classification.

Selected 2D classes were then subjected to 3D *ab initio* volume reconstruction as five classes. The dominant class with 846,965 particles was subjected to two rounds of heterogenous refinement against a junk class resulting in 838,538 particles, which refined to an overall resolution 3.96 Å and further improved to 3.43 Å upon non-uniform refinement. A generous mask around the complex was used in local refinement with a fulcrum at centre mass, which resulted in a 3.3 Å reconstruction. This map was sharpened using DeepEMhancer for representation(*35*).

#### Model refinement and structural analysis

Both sharpened and unsharpened maps from non-uniform refinement were used to build or fit models into the map. Chimera X (*36*) was used to dock individual ST2 molecules derived X-ray structures (PDB ID= 5VI4) into the density, whereas HpBARI2 was built manually using Coot (*37*). To improve model quality, Coot and phenix.realspace were combined iteratively to fix steric clashes, side chain and Ramachandran outliers (*38*). Interfaces were analysed using PISA, ChimeraX. All structural figures were made using ChimeraX.

### Surface plasmon resonance analysis (SPR)

#### Protein biotinylation and the experimental setup for SPR analysis

Avi-tagged ST2^ecto^ and IL-1RAcP^ecto^ were enzymatically biotinylated using the BirA500 Biotin Ligation Kit (Avidity LLC). Purified HpBARI2 or variants of HpBARI2 were chemically biotinylated by incubating 50 μM protein with 100 µM EZ-Link^TM^Sulfo-NHS-Biotin (Thermo Fisher Scientific) on ice for 2 h. Excess biotin was removed by buffer exchange using CentriPure 5 desalting columns (emp BIOTECH), which were washed and pre-equilibrated with 1× PBS. The biotinylated proteins were used as immobilized ligands for SPR analysis.

All SPR experiments were performed at 25°C on a Biacore T200 instrument (GE Healthcare) using SPR buffer (1x PBS + 0.05% Tween-20) and Biotin CAPture Kit (Cytiva). The sensor chip surface was first coated with streptavidin by injecting the CAPture reagent supplied with the kit at a flow rate of 2 µL/min for 300 s. Biotinylated proteins were then injected over the streptavidin-coated surface at 2 µL/min to immobilize the ligands at the desired response levels (RUs).

Binding interactions were assessed by injecting the individual binding partners over sensor surfaces with and without immobilized ligand at a flow rate of 30 µL/min. Following each binding cycle, the sensor surface was regenerated by injecting a mixture of 6 M guanidine-HCl and 1 M NaOH (pH 11.0) in a 3:1 ratio. The resulting sensograms were processed using BIAevaluation software version 1.0 (Biacore, GE Healthcare). Response curves were double referenced by subtracting the response obtained from the reference cell and the response from the corresponding blank injection.

#### Assessing binding of HpBARI2 and its active site variants to ST2^ecto^

To assess the impact HpBARI2 mutations on ST2 binding, single concentration SPR analysis was conducted and binding responses measured for each mutant relative to WT protein. A total of 500-600 response units (RU) of ST2^ecto^ was immobilised on the chip, and binding measurements were performed at a flow rate of 30 µl/min by injecting WT or mutant HpBARI2 at 5 nM concentration over the chip surface. Association was measured for 180 s, followed by dissociation for 240 s. After each binding cycle, the sensor chip surface was regenerated by injecting regeneration solution. The double referenced RU value at stability was used to calculate ST2 binding percentages as measures of binding perturbation of HpBARI2 mutants compared to the WT.

#### ST2^ecto^-HpBARI2 interaction kinetics

To obtain binding K_D_ for the interaction of ST2^ecto^ with WT or heterodimeric HpBARI2, 200- 300 RU of chemically biotinylated HpBARI2 proteins were captured on the chip surface.

Following ligand immobilization, a 2-fold dilution series of purified ST2^ecto^ (500 nM-15.625 nM) was flowed over the chip surface at a flow rate of 30 µl/min for 180s, followed by dissociation for 240 s. After each binding cycle the chip surface was regenerated as described in the experimental setup above.

The K_D_ for interaction between HpBARI2^mono^ and ST2^ecto^ was determined in two separate SPR formats. In the first format, 300-400 RU of ST2^ecto^ was immobilized followed by injection of 2-fold concentration series HpBARI2^mono^ (500 nM- 15.625 nM). In second format, the chemically biotinylated HpBARI2^mono^ was captured on the surface and a 2-fold concentration series of ST2^ecto^ (500 nM-15.625 nM) was flowed over the chip surface. In all cases, the K_D_ was calculated by fitting double reference subtracted response curves with 1:1 Langmuir model.

#### Analysis of IL-1RAcP^ecto^ binding

200-300 RU of monobiotinylated IL-1RAcP were immobilized on the chip surface. 100 nM of ST2^ecto^:IL-33 or ST2^ecto^:HpBARI2 complex were flowed over the chip surface for 120 s followed by dissociation measurement for another 120s.

#### ST2^ecto^ binding to ST2^ecto^:HpBARI2 complex

400-500 RU of monobiotinylated ST2^ecto^ were immobilized on the chip surface. HpBARI2 at 5 nM concentration was then injected over chip surface flow rate of 30 μl/min for 240s, followed immediately by injection of ST2^ecto^ at various concentrations ranging from 500nM- 1.25nM.

### Analysis of ST2-mediated signalling using *in vitro* bone marrow derived ILC2 cells

ILC2 reporter assays were used to directly compare suppressive effects of HpBARI2 and its variants on ST2:IL-33 signalling. Bone marrow single cell suspensions were prepared from 6- 10-week-old C57BL/6 J both male mice (Charles River), by flushing tibias and femurs with RPMI 1640 medium using a 21 g needle. Cells were resuspended in ACK lysis buffer (Gibco) for 5 min at room temperature, prior to resuspension in complete RPMI (with 10% FCS, 1% Penicillin/Streptomycin, 1% L-glutamine, Gibco) and passing through a 70 μm cell strainer.

Cells were cultured in round-bottom 96-well-plates in a final 200 μl volume, containing 0.5 × 10^6^ cells/well. IL-2 and IL-7 (Biolegend) were added at 10 ng/ml final concentration, IL- 33 (Biolegend) at 1 ng/ml final concentration. WT and its active site mutants were added in a range of protein concentrations. The suppressive effects of HpBARI2 were compared with those of HpBARI2^mono^ at protein concentrations corresponding to an equivalent ST2 binding sites, calculated based on molar weight per ST2 binding site (i.e. WT HpBARI2 = 59 kDa, with 3 ST2 binding sites = 19.7 kDa/binding site; HpBARI2^mono^ = 42 kDa, with 1 ST2 binding site = 42 kDa/binding site). Cells containing proteins were cultured at 37°C, 5% CO_2_ for 5 days, prior to collection of supernatants. IL-5 and IL-13 concentration were assessed in supernatants following manufacturer’s instructions using mouse uncoated IL-5 and IL-13 ELISA kits (Invitrogen).

### Analysis of HpBARI binding to ST2-containing liposomes

#### Preparation of ST2somes

ST2somes or liposomes bound ST2 were prepared using a lipid composition designed to approximate the plasma membrane of mouse immune cells based on published lipidomic data (*39*). All lipids were purchased from Avanti Polar Lipids and prepared as 20 mg/mL stock solutions in chloroform. Next, lipids were mixed in volume-to-volume ratio to generating a lipid mixture comprising 60% 1-palmitoyl-2-oleoyl-glycero-3-phosphocholine (POPC), 15% 1- palmitoyl-2-oleoyl-sn-glycero-3-phosphoethanolamine (POPE), 8% sphingomyelin, 15% cholesterol, and 2% 1-palmitoyl-2-oleoyl-sn-glycero-3-phospho-L-serine (POPS). This lipid mixture was dried under nitrogen stream in a fume hood, followed by vacuum drying for 2 h to remove residual chloroform. Dried lipids (10 mg) were resuspended in 500 µL of 1x PBS and fully solubilized with CHAPS at a final concentration of 5% (v/v). DDM-purified ST2 (1000 µl at 1.5 mg/mL) was added to the solubilized lipid mixture and incubated for 30 min at room temperature. The ST2-lipid mixture was further diluted to 3 ml using 1x PBS, and detergents were removed by dialysis for 48 h to promote incorporation of ST2 into the lipid bilayer. Following dialysis, the sample was centrifuged at 200,000 x g for 45 min using TLA 120.2 rotor (Beckman Coulter) to recover the lipid-associated ST2 fraction. The resulting white, amorphous pellets were resuspended in 3 ml of 1x PBS. SDS-PAGE analysis of resuspended lipid pellet and supernatant indicated that approximately 80% of the ST2 present following dialysis was recovered in the lipid-associated fraction. Next, the resuspended pellet centrifuged at 200,000 x g for 30 min. The pellet was resuspended in 0.8 ml of 1x PBS and this resuspension was extruded 25 times past a 100 nm filter (Avanti). The extruded suspension was then mixed with 3.2 mL of 60% (w/v) sucrose and overlaid sequentially with 3 mL of 30% (w/v) sucrose and 0.8 mL of 1x PBS. The samples were centrifuged at 200,000 x g for 2 h using an SW55Ti rotor (Beckman coulter). The upper fraction containing ST2somes was collected and dialyzed overnight against 1x PBS to remove residual sucrose. The dialyzed sample was centrifuged at 200,000 x g for 30 min, and the resulting ST2some pellet was resuspended in 1 mL of 1x PBS for subsequent experiments. ST2some size and homogeneity were assessed using dynamic light scattering. The total incorporated ST2 incorporated was determined by densitometric analysis of Coomassie-stained SDS-PAGE gels using DDM-purified ST2 of pre-determined concentration as a reference. Based on this analysis, 10 µl of ST2some suspension contained approximately 10 µg of ST2 protein. No appreciable ST2 aggregation was observed during the incorporation procedure.

#### ST2some binding and competition assays

For competition assays, 10 µl of ST2somes were first mixed with 190 µl 1xPBS. Assuming an approximately equal distribution of ST2 orientations across the two leaflets of the liposomal membrane, 50% of the ST2 molecules were assumed surface accessible, with a density of 1500-2000 ST2 molecules/µm^2^. Thus, 10 µL of ST2some suspension contained 5 µg of surface-accessible ST2. sST2 (2.9 µg; 37 kDa) was added to ST2some resuspension at equimolar quantities compared to surface-accessible membrane ST2 (62 kDa). HpBARI2 or HpBARI2^mono^ were added to final concentrations of 50 nM, and reaction mixtures were incubated for 30 min at 37 °C. Following incubation, reaction mixtures were centrifuged at 200,000 x g for 30 min to separate ST2somes from soluble proteins. The supernatant was collected, and the ST2some pellet was resuspended in 200 µl of 1x PBS. Aliquots of 5 µl from the resuspended pellet and supernatant fractions were analysed by SDS-PAGE and Western blotting. HPBARI2 and HpBARI2^mono^ were detected using an anti-cMyc antibody (Bio-Rad), whereas ST2 and sST2 were detected using an anti-His antibody (Qiagen). Empty liposomes (100 nm extruded) were used as controls for these binding experiments. Same experimental setup was used to assess the impact of sST2 concentration series (1.6µM-0.1 µM) on ST2- HpBARI2 interactions. Band densities on blots were estimated using Image J software.

### Circular Dichroism

Circular dichroism was performed with a Jasco J- 815 spectropolarimeter (Jasco, Japan). Spectra were measured in duplicates with 0.5 nm step resolution, a response time of 1 s and a scanning speed of 10 nm/min. The spectrum for each HpBARI2 and its variants was recorded from 190 to 250 nm at a protein concentration of 0.2 mg/ml in 0.1 mm quartz cuvette. The baseline (20 mM phosphate buffer pH 7.2 and 50 mM NaF) was subtracted from all measurements.

## Acknowledgments

Authors would like to thank Dr Rishi Matadeen at COSMIC cryoEM facility (University of Oxford) for help with data collection and processing. We thank David Staunton for support with biophysical measurements. We thank Hannah Ivison for laboratory management.

## Author contributions

A.J, H.J.M, and M.K.H conceived and planned the study and wrote the manuscript. A.J. conducted protein production, interaction analysis/biophysics studies and cryo-EM structure determination. NPO and SHH conducted ex vivo signalling assays. All authors designed experiments, analysed data, read and commented on the manuscript.

## Data availability

Data within graphs and uncropped gels and blot images are included with this submission as source data. Cryo-EM maps and models are deposited in the EM data bank and protein data bank with accession codes EMD-59759 and 34DU.

---

**Fig. S1.**
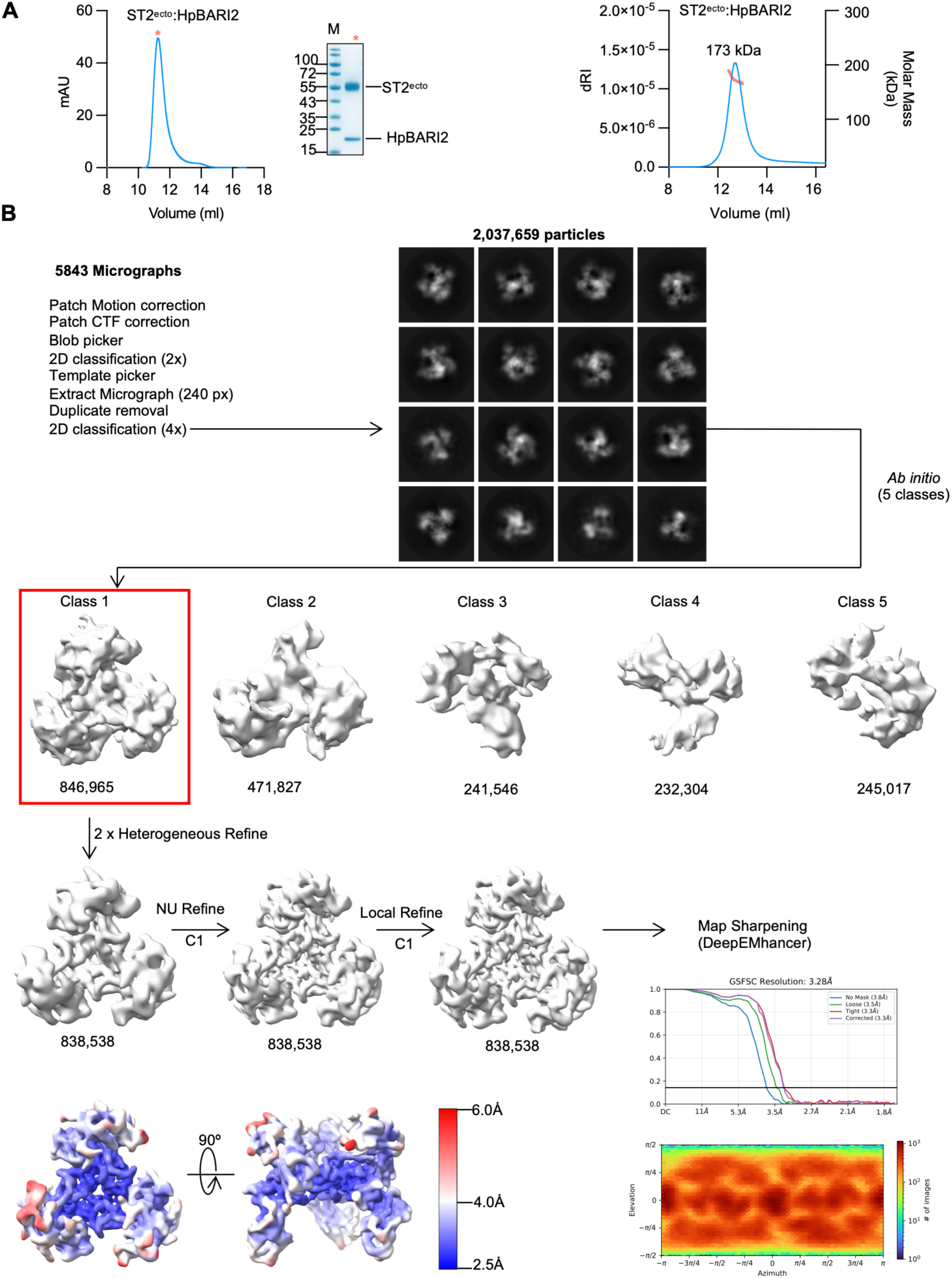
Determination of the cryo-EM structure of the ST2^ecto^:HpBARI2 complex. **(A)** Size exclusion chromatography trace showing purified ST2^ecto^:HpBARI2 complex (left panel). The y axis shows UV absorption at 280 nm in milli Absorbance Units (mAU) and the x axis shows the elution volume in millilitres. A red asterisk denotes the SEC peak fraction chosen for SDS-PAGE analysis by coomassie staining. Molecular weight estimation of ST2^ecto^:HpBARI2 by SEC-MALLS (right panel). The SEC trace in light blue shows absorbance at 280 nm while the coral-coloured line across the trace shows the molecular mass distribution of ST2^ecto^:HpBARI2 as determined by SEC-MALLS. The left y axis shows differential refractive index (dRI) and the right y axis depicts molar mass in kDa. The x axis show elution volume in millilitres (ml). **(B)** Workflow used to obtain high-resolution reconstructions of the ST2^ecto^:HpBARI2 complex. Particles from selected 2D classes were used to reconstruct *ab initio* 3D volumes. The dominant *ab initio* class shown in the red box was chosen for further refinement. All 3D volumes shown are unsharpened maps. In the bottom-right panel, the upper graph shows the gold-standard Fourier shell correlation (GSFSC) curve and corresponding resolution estimate, while the lower graph shows the angular distribution of particles used for 3D reconstruction and refinement.

**Fig. S2.**
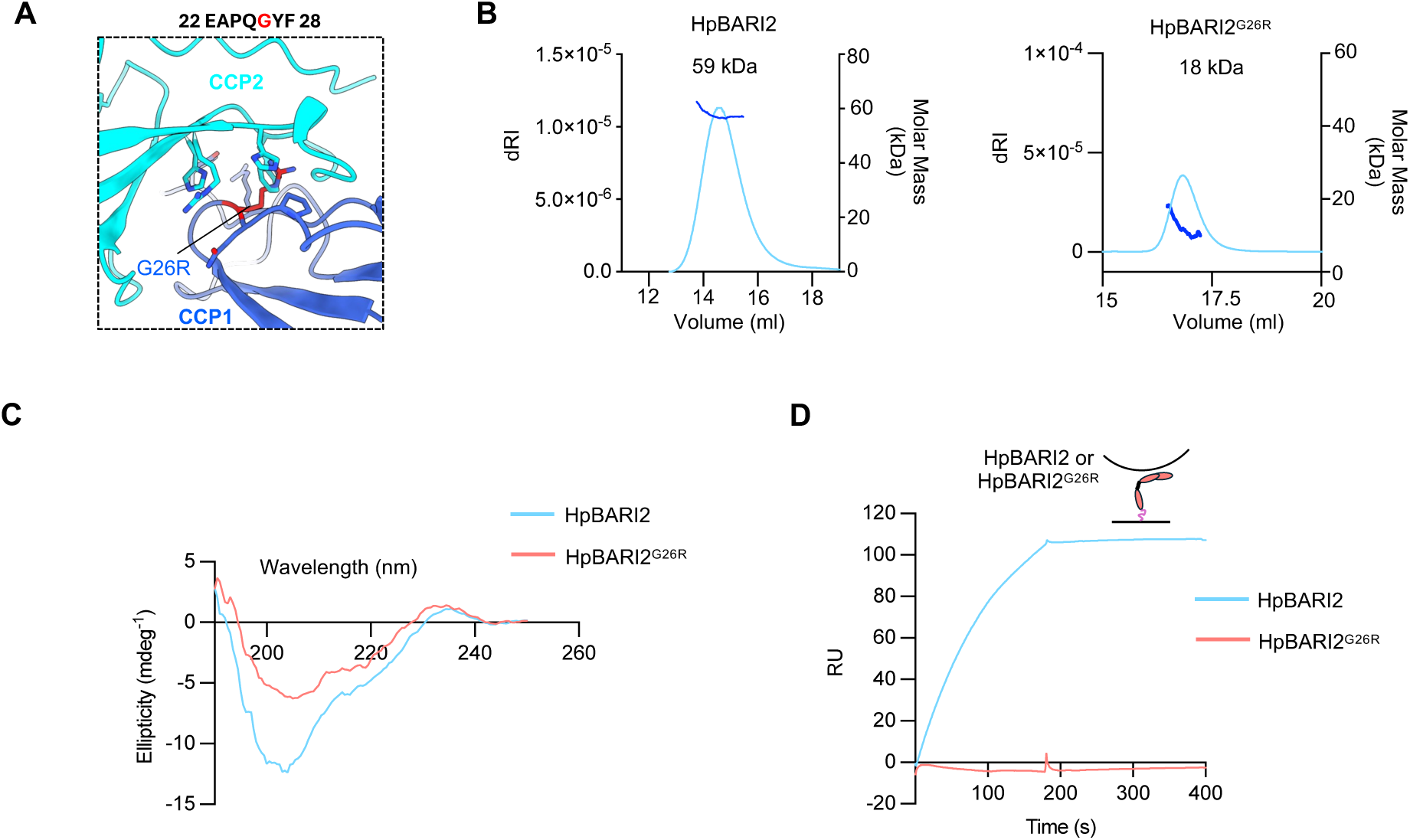
The trimeric arrangement of HpBARI2 is required for ST2 binding. **(A)** A zoomed-in view of the HpBARI2 vertex illustrating the CCP1:CCP2 packing interface. A G26R mutation is shown as red stick, and the native glycine residue within the HpBARI2 protein sequence is highlighted. G26R causes a steric clashes with surrounding side chains of residues from CCP2 shown as cyan sticks**. (B)** SEC-MALLS analysis of HpBARI2 (left) and HpBARI2^G26R^ (right). The light blue lines show absorbance at 280 nm while the dark blue lines show the molecular mass distribution of HpBARI2 and HpBARI2^G26R^ as determined by SEC-MALLS. The left y axis shows differential refractive index (dRI) and the right y axis depicts molar mass in kDa. The x axis show elution volume range for each protein in millilitres (ml). **(C)** Circular dichroism analysis of HpBARI2 and HpBARI2^G26R^. **(D)** SPR analysis showing binding of HpBARI2 and HpBARI2^G26R^ to ST2. 5 nM HpBARI2 or 500 nM HpBARI2^G26R^ were flowed over an ST2^ecto^ coated chip surface. This experiment was performed two times (n=2).

**Fig. S3.**
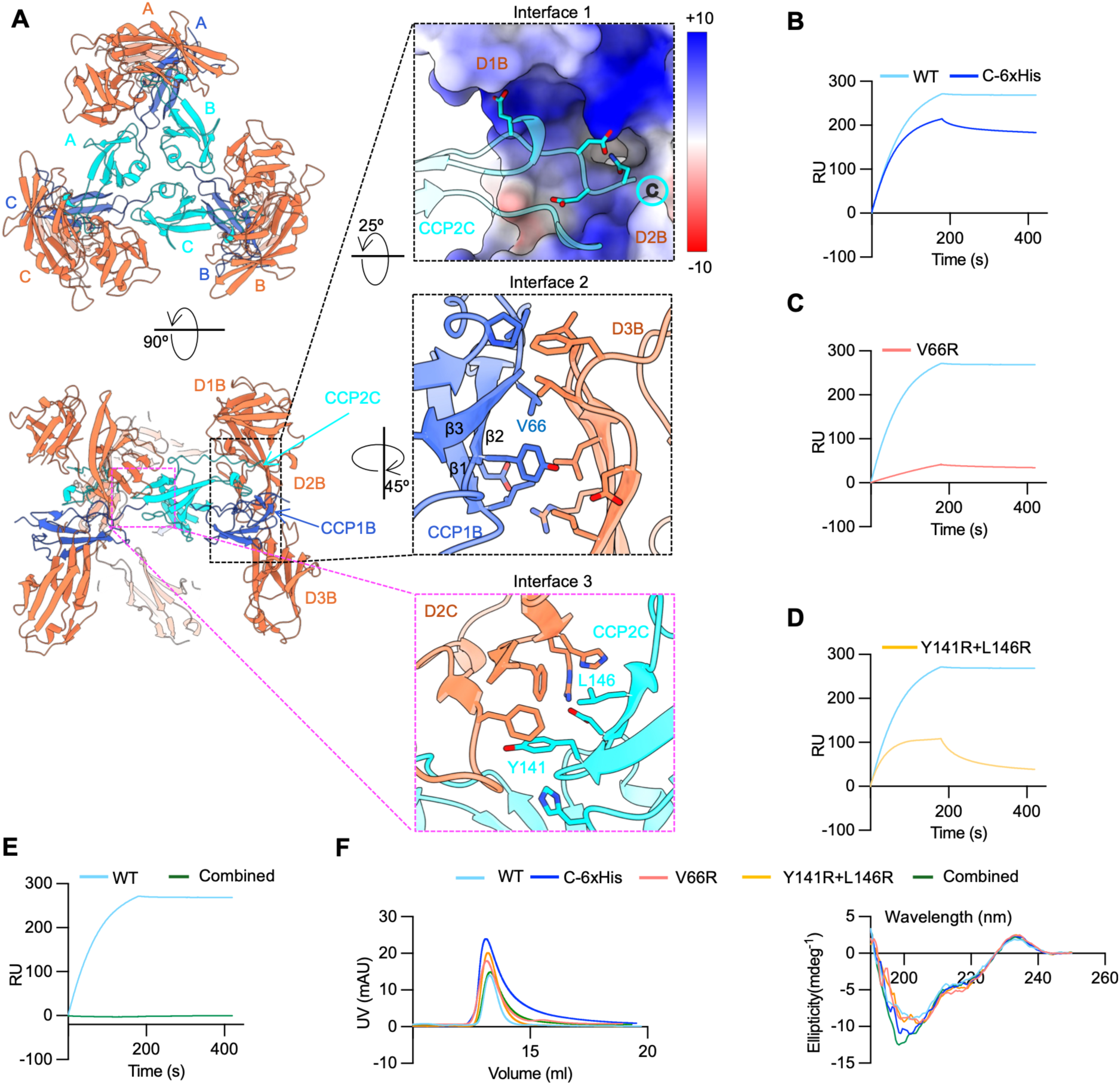
The ST2^ecto^:HpBARI2 complex contains three distinct interaction interfaces. **(A)** Two views of the ST2^ecto^:HpBARI2 complex depicted as cartoons. Enlarged views in dashed boxes show three key interfaces within the complex. Interfacial residues are shown as sticks. **(B–E)** SPR analysis of the interaction of wild type (WT) HpBARI2 and its interface mutants with ST2^ecto^ with the same WT curve shown in each case. HpBARI2 proteins were flowed over an ST2^ecto^-coated chip surface at a concentration of 5 nM, except for the combined mutant, which was analysed at 500 nM. **(F)** The left panel shows SEC chromatography traces for WT HpBARI2 and its interfacial variants. The right panel shows circular dichroism spectra of HpBARI2 and interface mutants.

**Fig. S4.**
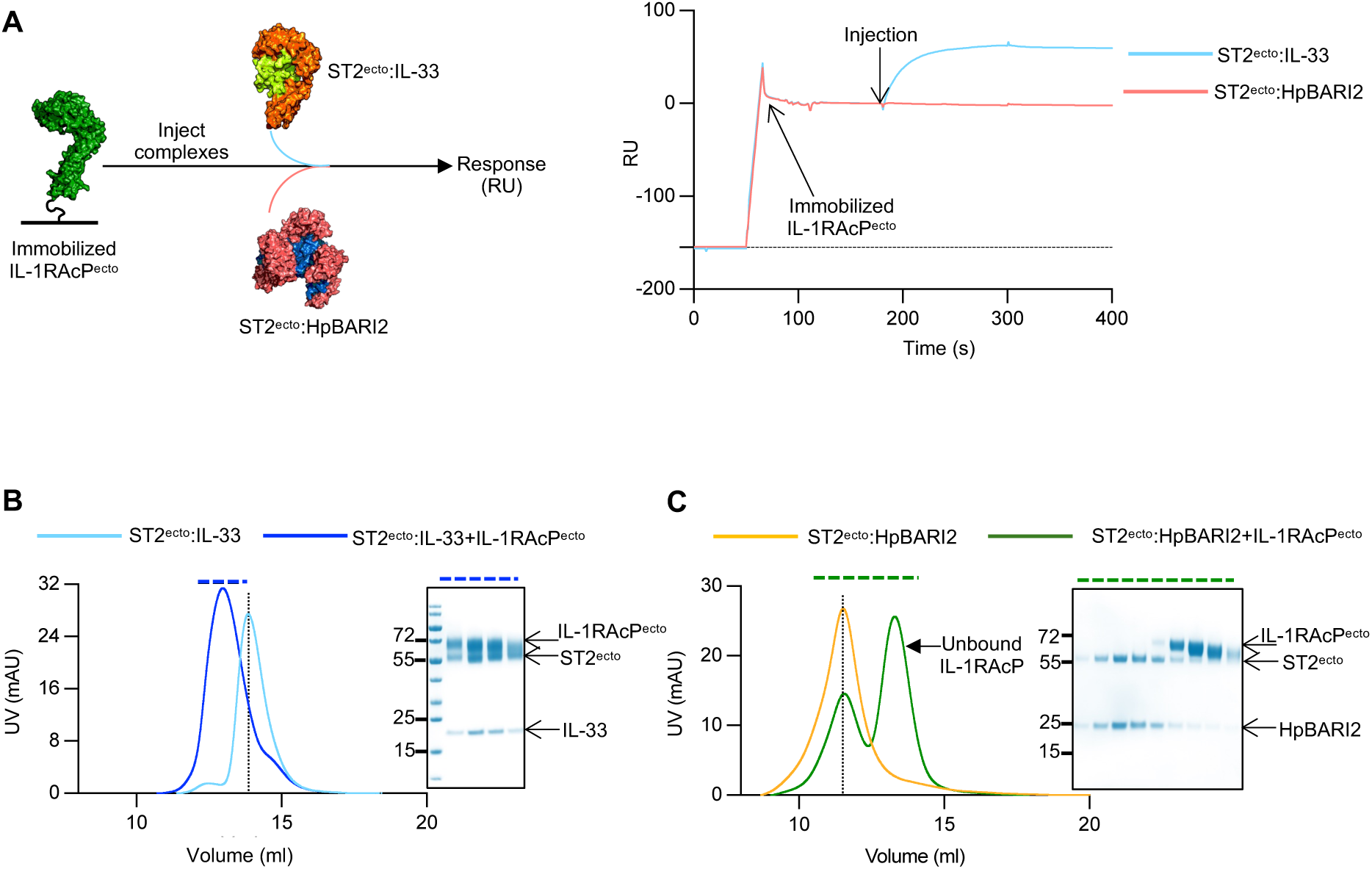
HpBARI2 binding prevents the interaction between ST2 and IL-1RAcP. **(A)** The left panel shows the SPR experimental setup used to assess IL-1RAcP^ecto^ binding to ST2^ecto^-containing complexes. The right panel shows representative real-time binding responses for interaction between ST2^ecto^:IL-33 and ST2^ecto^:HpBARI2 complexes with immobilised IL-1RAcP. The SPR sensograms are representative of two independent experiments. **(B)** SEC analysis of IL-1RAcP^ecto^ binding to ST2^ecto^:IL-33 complex. The left panel shows the SEC profiles of ST2^ecto^:IL-33 binary complex (light blue) and ST2^ecto^:IL-33:IL- 1RAcP^ecto^ ternary complex (dark blue). The right-hand panel shows Coomassie stained SDS-PAGE confirming co-fractionation of individual components from the ternary complex. The fractions analysed on the gel are indicated by a blue dashed line above the SEC trace. **(C)** SEC analysis of IL-1RAcP^ecto^ binding to ST2^ecto^:HpBARI2 complex. The left-hand panel shows the SEC profiles of the ST2^ecto^:HpBARI2 complex (orange) and the ST2^ecto^:HpBAR2:IL-1RAcP^ecto^ mixture (green). Addition of IL-1RAcP^ecto^ to ST2^ecto^:HpBARI2 does not shift the SEC profile (green) relative to the ST2^ecto^:HpBARI2 complex (orange) and unbound IL-1RAcP1 elutes as a distinct peak. The right-hand panel depicts, a Coomassie stained SDS-PAGE which further confirms that IL-1RAcP^ecto^ does not cofractionate with the ST2^ecto^:HpBARI2 complex. Fractions analysed on the gel are indicated by the dashed line above SEC trace (green). The SEC profiles represent two independent experiments (n=2).

**Fig. S5.**
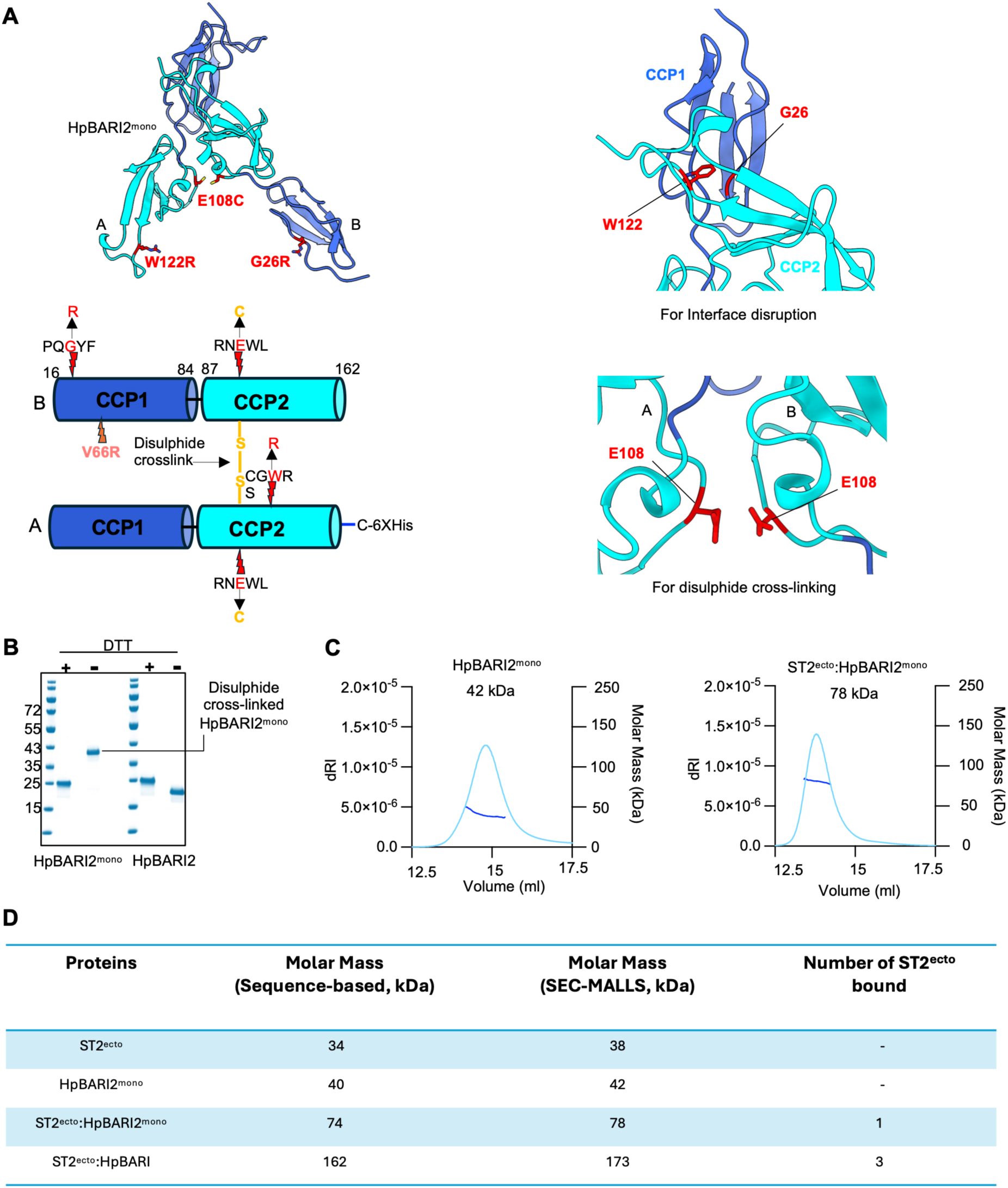
Design and characterization of monovalent HpBARI2. **(A)** Left, cartoon representation and domain schematic illustrating the design of HpBARI2^mono^. Residue substitutions introduced to disrupt trimer formation, enable disulphide cross-linking are shown as red sticks. Mutations or modifications that disrupt ST2 binding are marked on the domain schematic below. Right, cartoon views of the HpBARI2 trimer interfaces showing location of mutations. **(B)** Coomassie-stained SDS-PAGE gel showing the disulphide cross-linked HpBARI2^mono^. **(C)** SEC-MALLS analysis of HpBARI2^mono^ and its complex with ST2^ecto^. **(D)** Table showing the molecular weights and stoichiometries of ST2^ecto^ complexes with WT and HpBARI2^mono^, as inferred from the measured molecular weights.

**Fig. S6.**
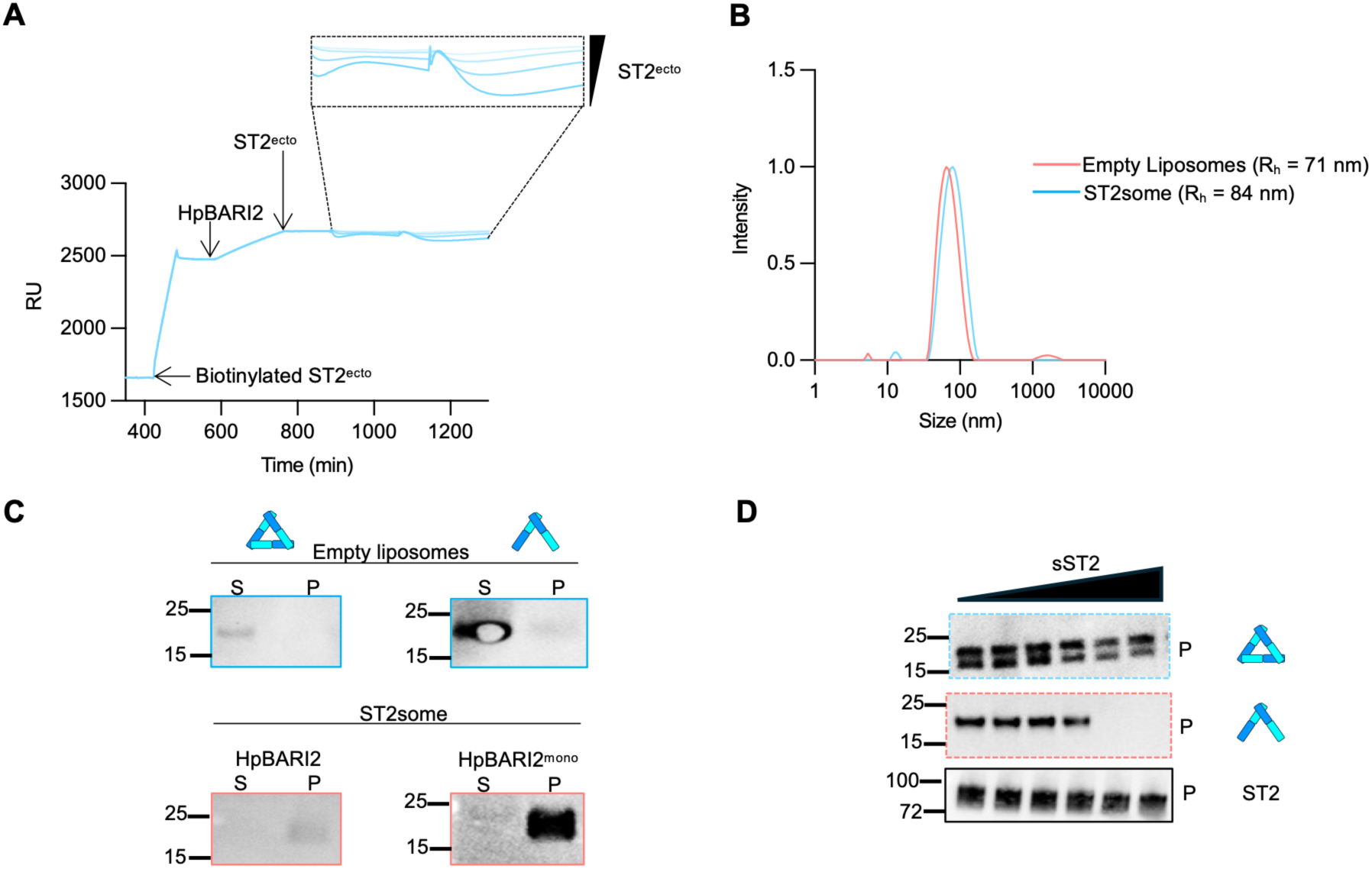
Characterization of HpBARI2 binding to ST2^ecto^ and ST2somes. **(A)** SPR sensograms showing little or no additional response upon flowing ST2^ecto^ over a chip surface coated with HpBARI2 pre-bound to immobilized ST2^ecto^. The enlarged view shows the response profiles obtained with the various concentrations of ST2^ecto^ (500-61.25 nM). **(B)** DLS analysis of extruded empty liposomes and ST2somes. **(C)** Western blot analysis of binding of HpBARI2 or HpBARI2^mono^ to empty liposomes and ST2somes. Both HpBARI2 and HpBARI2^mono^ pellet with ST2somes but not with empty liposomes. **(D)** The effect of a concentration series of sST2 on the interaction of HpBARI2 or HpBARI2^mono^ with ST2somes. One representative of two biological replicates is shown.

**Table 1.** Cryo-EM data collection, refinement and validation statistics for the ST2^ecto^:HpBARI2 complex.

| **Data collection and processing** | **EMDB: EMD-59759**<br>**PDB: 34DU** |
| --- | --- |
| Microscope | Titan Krios |
| Detector | Gatan K3 with 20 eV filter |
| Magnification | 58149 |
| Voltage (kV) | 300 |
| Electron exposure (e/Å<sup>2</sup>) | 48.97 |
| Defocus range (μm) | -3 to -1 |
| Pixel size (Å) | 0.832 |
| Symmetry | C1 |
| Number of particles | 838,538 |
| Map resolution (Å) | 3.3 |
| FSC threshold | 0.143 |
| Map resolution range (Å) | 3.3-4.0 |
| FSC threshold | 0.5 |
| **Refinement** |  |
| Initial model used | 5VI4 |
| Model resolution | 3.37 |
| FSC threshold | 0.5 |
| **Model Composition** |  |
| Non hydrogen atoms | 10572 |
| Amino acid residues | 1329 |
| Ligands | NA |
| **B-factor (Å<sup>2</sup>)** |  |
| Protein | 102.66 |
| Ligand | NA |
| **RMSD** |  |
| Bond length (Å) | 0.003 |
| Bond angles (°) | 0.595 |
| **Validation** |  |
| Molprobity score | 1.66 |
| Clash score | 8.74 |
| Rotamer outliers (%) | 2 |
| **Ramachandran plot** |  |
| Favoured (%) | 97.0 |
| Allowed (%) | 3.0 |
| Outliers (%) | 0 |
| **Model vs data fit** |  |
| CC<sub>mask</sub> | 0.86 |
| CC<sub>box</sub> | 0.80 |

**Table 2.** Description of interactions between ST2^ecto^ with HpBARI2.

| *Mus musculus* ST2<sup>ecto</sup> |  |  | HpBARI2 |  |  | Interaction |
| --- | --- | --- | --- | --- | --- | --- |
| D1-D2 |  |  | CCP2 |  |  |  |
| Chain | Residue | Group | Chain | Residue | Group |  |
| D/E/F | Lys136 | Side chain | A/B/C | Asp142 | Side chain | Hydrogen bond |
| D/E/F | Arg166 | Side chain | A/B/C | Pro147 | Backbone CO | Hydrogen bond |
| D/E/F | Arg169 | Side chain | A/B/C | Asp144 | Backbone CO | Hydrogen bond |
| D/E/F | Arg41 | Side chain | A/B/C | Asp 160 | Side chain | Electrostatic |
| D/E/F | Phe135 | Side chain |  |  |  | Hydrophobic |
| D/E/F | Phe173 | Side chain |  |  |  | Hydrophobic |
|  |  |  | A/B/C | Leu 146 | Side chain | Hydrophobic |
| D3 |  |  | CCP1 |  |  |  |
| D/E/F | Arg318 | Side chain | A/BC | Glu32 | Side chain | Hydrogen bond |
| D/E/F | Asp253 | Side chain | A/B/C | Tyr49 | Side chain | Hydrogen bond |
| D/E/F | Leu251 | Side chain |  |  |  | Hydrophobic |
| D/E/F | Phe250 | Side chain |  |  |  | Hydrophobic |
| D/E/F | Leu255 | Side chain |  |  |  | Hydrophobic |
| D/E/F | Leu309 | Side chain |  |  |  | Hydrophobic |
| D/E/F | Leu311 | Side chain |  |  |  | Hydrophobic |
| D/E/F | Met316 | Side chain |  |  |  | Hydrophobic |

